# Evaluating performance of metagenomic characterization algorithms using *in silico* datasets generated with FASTQSim

**DOI:** 10.1101/046532

**Authors:** Anna Shcherbina, Darrell O. Ricke, Nelson Chiu

**Author notes:** Corresponding author: Darrell Ricke Anna Shcherbina - Nelson Chiu.

## Abstract

**Background:** *In silico* bacterial, viral, and human truth datasets were generated to evaluate available metagenomics algorithms. Sequenced datasets include background organisms, creating ambiguity in the true source organism for each read. Bacterial and viral datasets were created with even and staggered coverage to evaluate organism identification, read mapping, and gene identification capabilities of available algorithms. These truth datasets are provided as a resource for the development and refinement of metagenomic algorithms. Algorithm performance on these truth datasets can inform decision makers on strengths and weaknesses of available algorithms and how the results may be best leveraged for bacterial and viral organism identification and characterization.

Source organisms were selected to mirror communities described in the Human Microbiome Project as well as the emerging pathogens listed by the National Institute of Allergy and Infectious Diseases. The six *in silico* datasets were used to evaluate the performance of six leading metagenomics algorithms: MetaScope, Kraken, LMAT, MetaPhlAn, MetaCV, and MetaPhyler.

**Results:** Algorithms were evaluated on runtime, true positive organisms identified to the genus and species levels, false positive organisms identified to genus and species level, read mapping, relative abundance estimation, and gene calling. No algorithm out performed the others in all categories, and the algorithm or algorithms of choice strongly depends on analysis goals. MetaPhlAn excels for bacteria and LMAT for viruses. The algorithms were ranked by overall performance using a normalized weighted sum of the above metrics, and MetaScope emerged as the overall winner, followed by Kraken and LMAT.

**Conclusions:** Simulated FASTQ datasets with well-characterized truth data about microbial community composition reveal numerous insights about the relative strengths and weaknesses of the metagenomics algorithms evaluated. The simulated datasets are available to download from the Sequence Read Archive (SRP062063).

## Background

Continuing advances in sequencing technologies are increasing the feasibility of sequencing entire microbial communities rather than individual organisms. This has led to rapid developments in the field of metagenomics aimed at studying genomic material recovered directly from environmental and medical samples. Sequencing the metagenome enables the capture of greater genetic diversity than can be sampled with highly targeted approaches such as microarrays. Metagenomic sequencing has a number of applications for medical diagnostics (i.e. human gut microbiome analysis), environmental profiling (i.e. soil samples), and homeland defense[1-3]. Metagenomic techniques also enable the study of communities of organisms simulated *in vitro*[4].

Simultaneously, a number of bioinformatics tools have been developed to analyze metagenomic sample data. They employ a variety of techniques to achieve the opposing goals of high accuracy and low runtime. In this study, the performance of these varied approaches to metagenomic sequence classification was evaluated on a suite of *in silico* datasets with perfectly characterized composition. MetaScope, winner of the Defense Threat Reduction Agency’s Grand Challenge[5], relies on sequence analysis using spaced seeds followed by an augmented least common ancestor algorithm to map reads and assign genes for input FASTQ samples[6, 7]. Kraken[8] uses exact alignment of k-mers in combination with an optimized database and another version of the least common ancestors algorithm. MetaPhlAn[9] relies on unique clade-specific marker genes identified from 3000 reference genomes. The Livermore Metagenomic Analysis Toolkit (LMAT) exploits genetic relationships between different organisms by pre-computing the occurrence of each short sequence across the entire reference database and storing the evolutionarily conserved sequence patterns[10-12]. MetaCV translates nucleotide sequences into six frame peptides, which are then decomposed into k-mers. The k-mer frequency is computed in a protein-reference database and used to assign k-mer weights[13]. Finally, MetaPhyler uses a precomputed database of reference phylogenetic marker genes to build a sequence classifier. The classifier, based on BLAST, uses trained thresholds for various combinations of taxonomic ranks, sequence length, and reference genomes[14].

Simulated *in silico* datasets are a valuable tool for metagenomic research and provide capabilities to evaluate algorithm performance as well as to test hypotheses that cannot be examined through empirical observation. For example, simulated data has revealed biases and heterogeneity in the estimation of diversity metrics from metagenomics samples[15]. Additionally, multiple studies have demonstrated the usefulness of simulated metagenomics datasets for benchmarking sequence assembly and gene prediction pipelines[16-18]. Simulated datasets are also an effective means of parameter optimization for improved algorithm performance and can be used to optimize study design. Sequence simulation can aid with answering questions about coverage requirements, necessary sequence length, and whether paired-end or single-end sequencing should be used. For example, the ART simulator was successfully used by the 1000 Genomes Project Consortium to examine the effects of read length and PE insert size on a read’s ability to map to the human genome[19].

In this study, six *in silico* datasets were simulated by the FASTQsim tool. **Figure S1** illustrates the composition of each dataset. These datasets contained sequences from reference bacterial and viral genomes, as most human pathogens are members of these taxa. The HMP Even and HMP Staggered datasets were generated to include sequences from the 20 organisms from the Human Microbiome Project[20] (**Supplementary Table 1**). The HMP organisms were selected for inclusion after an attempt to benchmark the performance of MetaScope with the HMP dataset revealed potential contamination in the dataset. As the HMP benchmark dataset was generated by sequencing organisms cultured *in vitro*, there was no absolute truth for any background contaminant organisms in the dataset and it was not possible to determine whether the contamination was real or whether MetaScope was calling false positive organisms.

The bacterial dataset (**Supplementary Table 2**) was designed to test algorithm specificity. Four genera of pathogens were selected from the National Institute of Allergy and Infectious Diseases list of biodefense and emerging infectious disease agents[21] due to their relevance to disease diagnostics from metagenomics samples. These included *Yersinia, Coxiella, Brucella*, and *Salmonella*. Additionally, the *Escherichia* genus was added to the list due to the high abundance of representative sequences in GenBank[22].

Two virus datasets were generated with 21 species across 11 representative genera (**Supplementary Table 3**). As with the bacterial dataset, candidates were selected due to their inclusion on the NIAID list of emerging pathogens (Marburg virus, Machupo virus, Sudan ebolavirus, Junin virus, Guanarito virus, Chapare virus, Omsk hemorrhagic fever virus) as well as abundance of representative organisms in GenBank (HIV 1, HIV 2, Influenza A virus).

Finally, a dataset of human reads from build GRCh38 at 10x (22 million reads) coverage was generated to test host-filtering capabilities of each algorithm. This dataset was generated to measure how well algorithms can overcome the challenges posed by human sequence contamination in public reference databases[23]. For example, endogenous retroviral remnants may be incorrectly classified as belonging to viral genomes in a sample[24-26].

## Methods

### Improvements to FASTQsim

The FASTQsim toolkit was augmented to annotate gene information for simulated reads[27]. The “FASTQmapGenes” functionality was added, allowing users to specify NCBI accession ids to use for annotating gene information in simulated reads. The FASTQsim toolkit uses the Entrez and SeqIO libraries from BioPython[28] to download the specified files from GenBank in .gb format. The GenbankParser[29] java application is then used to parse the .gb files in order to extract all information encoded in the CDS and Gene tags. These gene and CDS annotations are appended to the headers within the simulated FASTQ files generated by FASTQsim, such that all reads that fall within a CDS or gene region are annotated with the corresponding CDS and gene information.

### *In silico* data generation

The FASTQsim toolkit was used to generate six *in silico* datasets. All were generated with the Illumina error and read length profile included with FASTQsim version 2.0, with no host background added. Specifically, read length of 150 bases was used, with single base mutation, insertion, and deletion rates as specified in the FASTQsim v. 2.0 documentation (http://sourceforge.net/p/fastqsim/code/ci/master/tree/params/illumina/). NCBI identifiers for all input data are listed in **Supplementary Tables 1-3**. The Krona toolkit[30] was used to visualize evaluation dataset composition.

Two *in silico* datasets were generated – “HMP Even” and “HMP Staggered” (**Supplementary Table 1**). For the HMP even dataset, FASTQsim was executed to provide equal number of reads for each species of organism (approximately 60,000 reads per species), with one exception - 559 reads for *Streptococcus agalactiae* were added to simulate a low-level contaminant organism. Version 2.0 of the FASTQsim algorithm probabilistically simulated read counts and error distributions based on a provided model. Due to the probabilistic nature of the algorithm, coverage levels deviated slightly from the specified 60,000 reads, with the largest deviation observed for the E. faecalis organism (52,290 reads). For the HMP Staggered dataset, coverage levels varied from 11.3x (217,512 reads) for *Actinomyces odontolyticus* to 0.001x (2 reads) for *Neisseria meningitidis*. The goal of the staggered dataset was to evaluate the ability of metagenomic algorithms to detect organisms present at very low concentrations, i.e. less than 5 reads.

The bacterial dataset included reads from the genear *Yersinia, Coxiella, Brucella, Salmonella*, and *Escherichia*. For each of the five genera, several representative species were selected (i.e., *Brucella abortus, Brucella melitensis, Brucella suis*). Next, several representative strains were selected for each species (i.e. *Brucella melitensis* ATCC 23457, *Brucella melitensis* biovar abortus 2308, *Brucella melitensis* biovar 1 strain 16M, and *Brucella melitensis* M28). Organisms were spiked into a FASTQ dataset with coverage levels ranging from 10x to 0.00002x (1 read).

For the Virus Even dataset, 10x coverage of each organism was simulated. For the Virus Staggered dataset, coverage varied from 100x for Sudan ebolavirus to 0.5x for the Human coronavirus HKU1.

### Metagenomic algorithm execution

Six metagenomic algorithms were selected for execution on the evaluation datasets. These included:

- MetaScope – winner of the Defense Threat Reduction Agency’s Grand Challenge[7] (version 2.0)
- MetaPhlAn[9] (version 1.7.8, https://bitbucket.org/nsegata/MetaPhlAn/src/),
- MetaCV[13] (version 2.3.0, http://sourceforge.net/projects/metacv/files/),
- MetaPhyler[14] (version 1.13, http://MetaPhyler.cbcb.umd.edu/#download),
- Kraken[8] (v0.10.5, https://ccb.jhu.edu/software/kraken/),
- LMAT[10-12] (v1.2.5, http://sourceforge.net/projects/lmat/).

All algorithms were executed on each of the evaluation datasets using a machine with 512 GB of RAM, 64 cores, 1 TB hard drive, running the Fedora 17 operating system. All algorithms were executed with the default set of databases described in their respective documentation, downloaded on March 1, 2015. Algorithms were evaluated using 60 of the 64 available cores.

Attempts were also made to install and run the SURPI (v1.0, https://github.com/chiulab/surpi)[31] and compressed BLAST (v0.9, http://cast.csail.mit.edu/)[32] algorithms, but these were unsuccessful.

### Algorithm performance evaluation

Runtime in seconds, true positive genus and species calls, false positive genus and species calls, read mapping, and relative abundance results at the species level were computed for all algorithm results. Additionally, correct gene calls were calculated for the set of algorithms that provided gene calling results (MetaScope, MetaCV, LMAT). The Gene ID Conversion function in the DAVID Bioinformatics Database[33] was used to convert across gene representation formats utilized by the three algorithms. Genes were marked as true positives if they matched the gene id, official gene symbol, locus tag, protein id, or specific product name of the truth data.

## Availability of supporting data

The FASTQsim toolkit can be downloaded from SourceForge: http://sourceforge.net/projects/fastqsim/

*In silico* evaluation datasets can be downloaded from the Sequence Read Archive: SRP062063

SRR2146185 – Virus Staggered dataset

SRR2146184- Virus Even dataset

SRR2146183 - Bacterial dataset

SRR2146181**—**HMP Staggered dataset

SRR2146182 - HMP Even dataset

## Results and Discussion

Runtime in seconds, true positive genus and species identification, false positive genus and species identification, and false negative species calls were determined for each of the metagenomic algorithms (**Figure 1**). Among the algorithms evaluated, only MetaScope mapped a small number of reads in our datasets to a taxon rank below species. Consequently, although the initial focus of the Bacterial dataset was to assess the ability of the algorithms to distinguish between different strains of the same species, it was decided to evaluate both true and false positives at species and genus level. To determine an overall rank of the algorithms across the datasets, the area occupied by each in the radar plot was computed (**Table 1**). When the polygon area was calculated using the MATLAB polyarea function and summed across all datasets, MetaScope emerges as the winner, with the largest overall area. Kraken and LMAT are the runner-ups, and MetaPhyler performed the worst. In addition to the algorithms’ rank overall, several trends can be noted in the individual performance categories.

**Figure 1.**
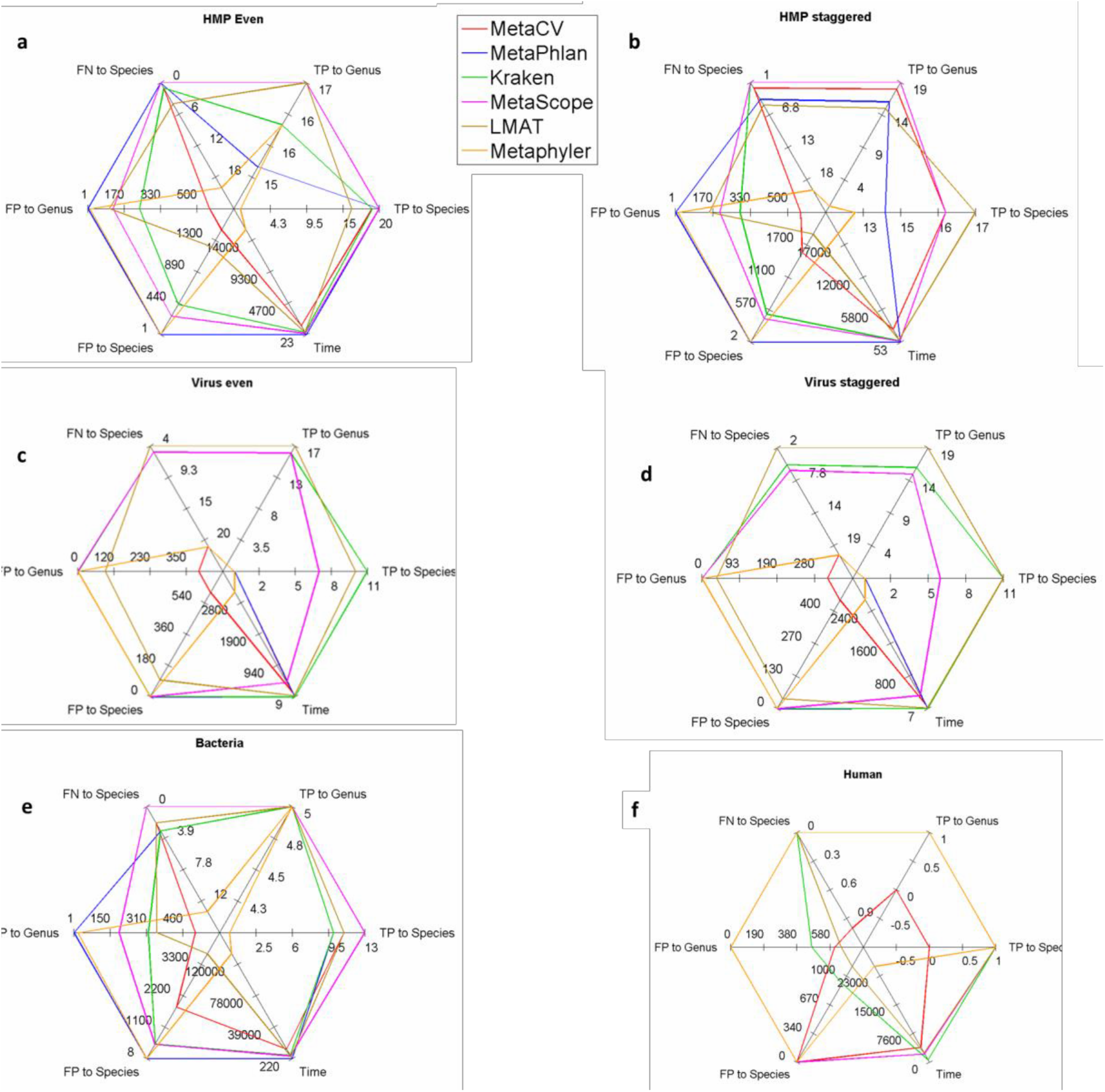
Performance metrics for 6 metagenomic analysis algorithms across the 6 in silico evaluation datasets. Algorithms evaluated include MetaCV (red line), MetaPhlAn (blue line), Kraken (green line), MetaScope (pink line), LMAT (brown line), MetaPhyler (orange line). Metrics evaluated include true positives (TP) to genus level, TP to species level, false positives (FP) to genus level, FP to species level, false negatives (FN) to species level, and runtime in seconds. Values indicative of high performance are at the periphery of the Radar plot, values indicative of poor performance are at the center of the plot. **a**. HMP dataset with even coverage. **b**. HMP dataset with staggered coverage. **c**. Virus dataset with even coverage. **d**. Virus dataset with staggered coverage. **e**. Bacterial dataset. **f**. Human dataset. The MetaPhlAn algorithm failed to run on the human dataset.

**Table 1.** Radar plot area in normalized units across six evaluation datasets. Higher areas, indicative of better performance, are colored in blue.

| Dataset | Human | Virus Staggered | Virus Even | Bacteria | HMP Staggered | HMP Even | Area Sum |
| --- | --- | --- | --- | --- | --- | --- | --- |
| MetaScope | 2.54 | 1.90 | 2.14 | 2.23 | 2.03 | 2.32 | 13.15 |
| Kraken | 1.64 | 2.36 | 1.21 | 1.75 | 1.90 | 1.76 | 10.62 |
| LMAT | 1.41 | 2.45 | 2.25 | 1.38 | 1.46 | 1.59 | 10.54 |
| MetaPhlAn |  | 0.48 | 0.99 | 2.25 | 1.88 | 2.02 | 7.62 |
| MetaCV | 0.82 | 0.09 | 0.14 | 1.43 | 1.25 | 1.08 | 4.80 |
| MetaPhyler | 1.88 | 0.60 | 0.09 | 0.67 | 0.57 | 0.63 | 4.44 |

The algorithms diverged in runtime by several orders of magnitude (**Table 2)**. Overall, MetaPhlAn had the shortest runtime. The algorithm had the fastest time on the three bacterial datasets – 22.64 s for HMP Even, 53.3 s on HMP staggered, and 220 s. on Bacteria. The second fastest times for these three datasets were 5 to 10 times slower: 233 s (MetaPhlAn), 261 s (MetaScope), and 2,700 s (LMAT), respectively. MetaPhlAn is able to execute quickly partly because it does not perform a host-filtering step. MetaPhlAn came in second for the virus datasets, with a runtime of 11 seconds on both, compared to 9 and 7 seconds for Kraken. MetaPhlAn failed to run on the human dataset. Kraken, MetaScope, and LMAT exhibited similar runtimes on all datasets, averaging 353 s on HMP Even, 354 s on HMP staggered, and 3,595 s on Bacteria. On the other end of the spectrum, MetaPhyler was an outlier for high runtime, requiring 15,480 s on HMP Even, 19,231 s on HMP staggered, and 129,600 s on Bacteria.

**Table 2.**
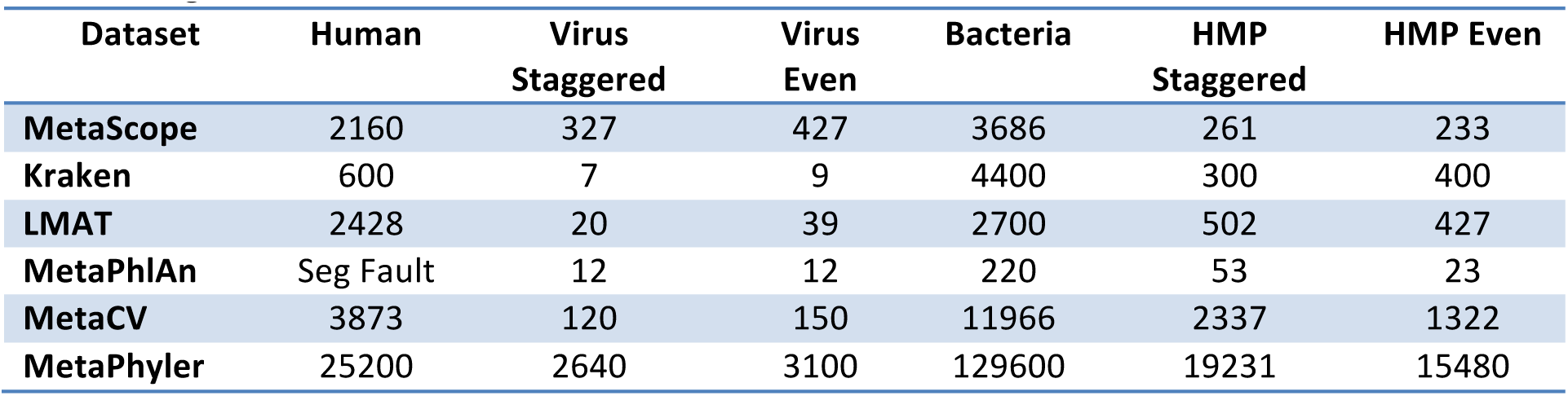
Algorithm runtime in seconds across six evaluation datasets.

In addition to its high speed, MetaPhlAn also achieved the highest accuracy, defined as ratio of true positives to false positives, on the bacterial datasets. It identified all 20 species in the HMP even dataset with only a single false positive organism. On HMP staggered, it missed 4 species out of 20 but reported only 2 false positive species. MetaScope, the runner up, reported a single false negative species but 414 false positives. However, the MetaPhlAn reference database is customized for bacteria, and no support exists at the time of this writing for profiling viruses or eukaryotes. MetaScope achieved the second- highest ratio of true positives to false positives, reporting slightly more true positives and approximately half as many false positives as Kraken. LMAT was the least conservative and reported the highest number of false positive organisms. MetaPhyler made highly conservative calls—false positives were low, but so were true positives. Additionally, MetaPhyler, and MetaCV, as well as MetaPhlAn, did not report results for the viral datasets.

Algorithm performance on the Human dataset (**Figure 2k**) illustrates the efficacy of the host-filtering step for each algorithm. The human reference genome is incomplete[34, 35] and misses regions specific to individual host subjects. These missed regions show up as false positives on the Human evaluation dataset – algorithms assign them to organisms other than the human host because these reads are not removed during the host filtering step. For example, MetaScope reports 152 organisms, with fewer than 100 reads assigned to each. Kraken has a similar false positive profile; it reports 1,266 species that account for <1% of the reads in the dataset. MetaCV reports 2,998 false organisms with low read count, and LMAT reports 1,118 species that account for less than 0.01% of the reads. MetaPhyler does not report results more specific than the Class taxonomy level for the Human dataset, in line with the conservative approach of this algorithm. MetaPhlAn crashes with a segmentation fault on the Human dataset, which most likely is an artifact of the non-host-filtering approach used by this algorithm.

**Figure 2.**
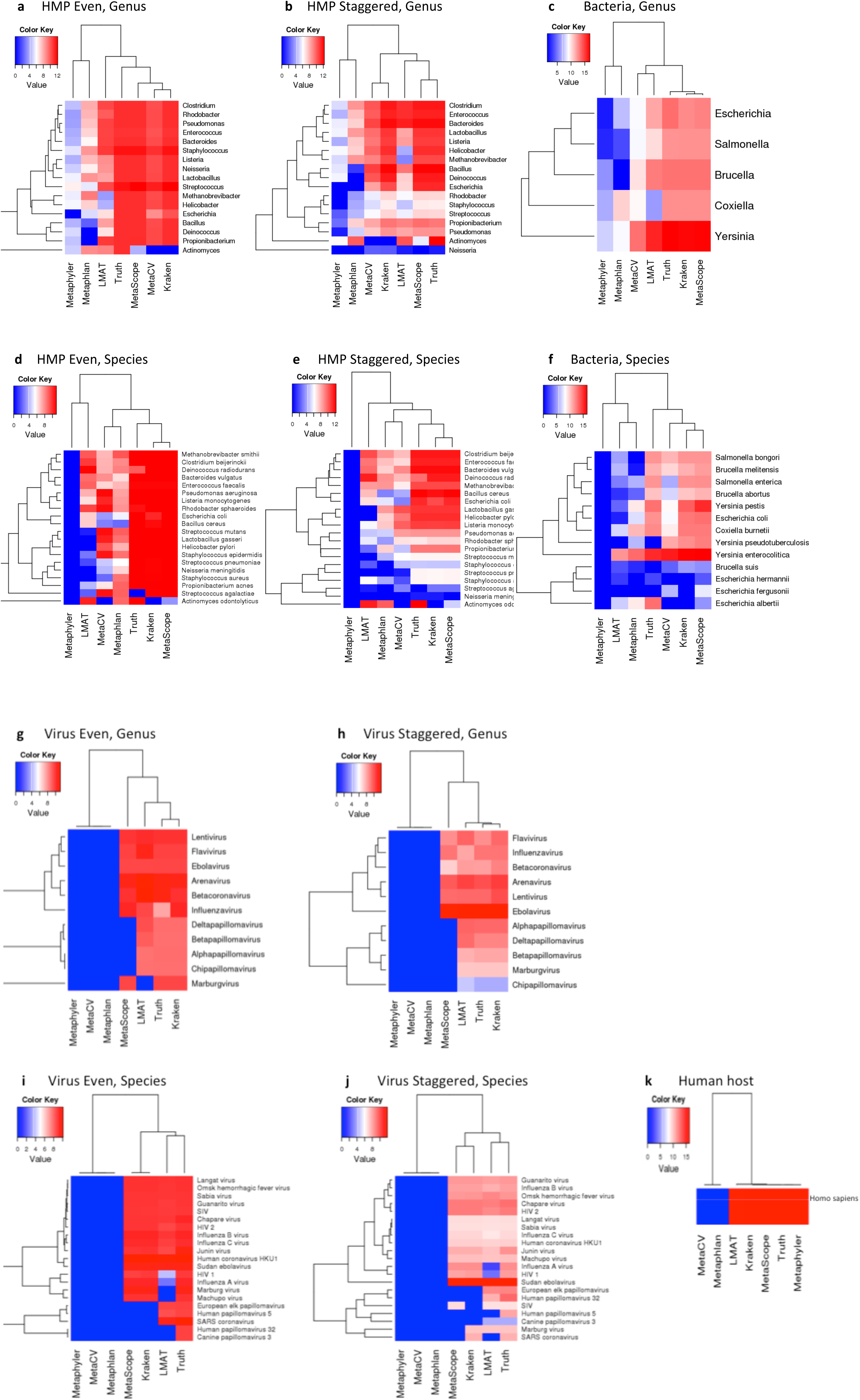
Number of correctly assigned reads to each organism at the genus and species level. Heatmap color scales are log10 (number of correctly assigned reads). The “Truth” column indicates the number of reads spiked into the FASTQ input file for the specified genus or species. **a.** – **e.** Reads mapped correctly to the genus level for the HMP even, HMP staggered, bacteria, virus even, virus staggered datasets, respectively. **f.** – **k.** Reads mapped correctly to the species level for HMP even, HMP staggered, bacteria, virus even, virus staggered, and human datasets, respectively.

The algorithms were evaluated based on their ability to correctly map reads and predict relative abundance of the organisms in the data (Figures 2, 3). For the bacterial datasets, Kraken and MetaScope classified the highest number of reads correctly for both the genus and species level, and cluster closest to the truth in the dendrogram. However, for the viral datasets, LMAT performed best, classifying the most reads correctly.

**Figure 3.**
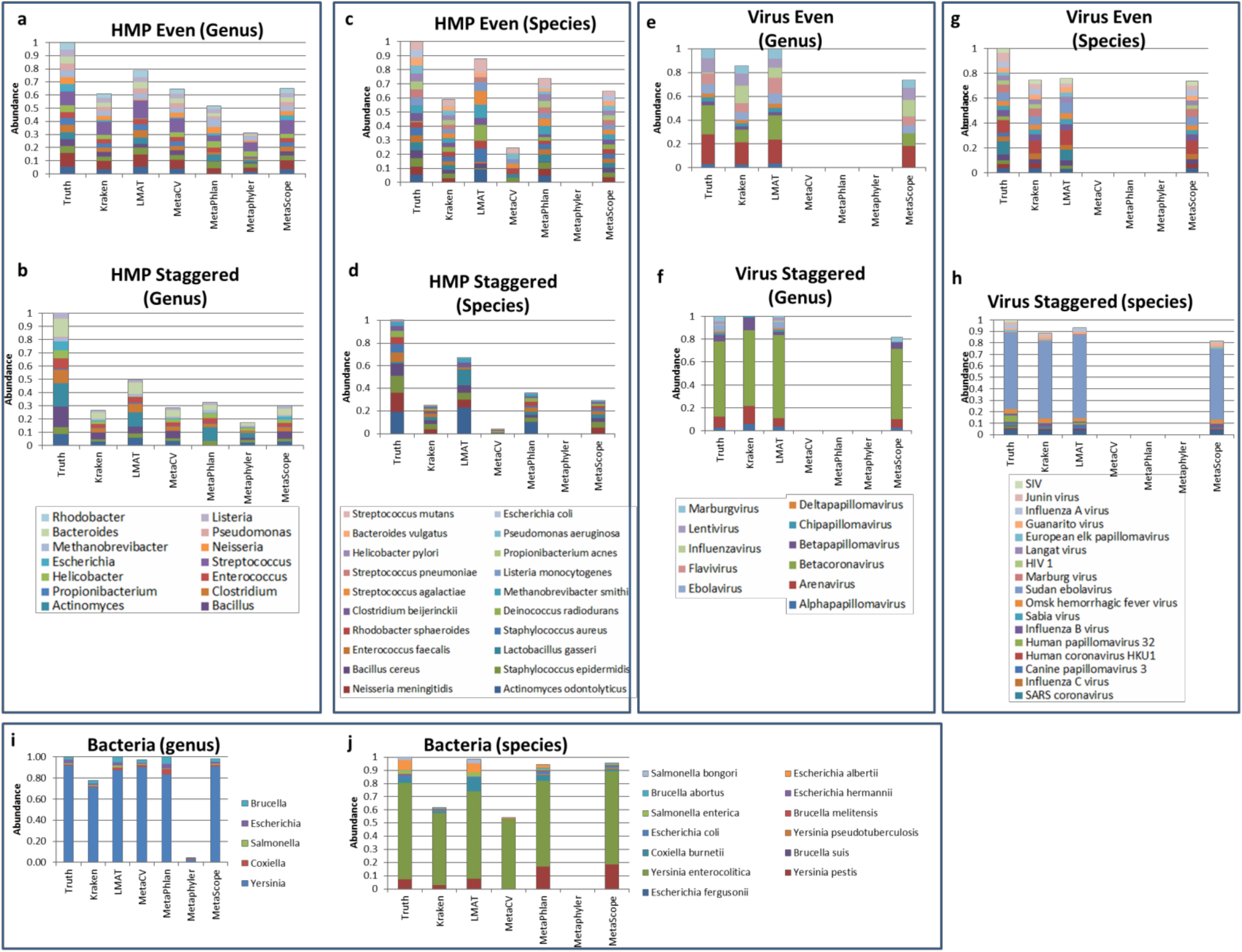
Relative abundance of organisms to the species and genus level. “Truth” column indicates relative abundance of genera and species added to the in silico FASTQ input file.

Although the *Actinomyces odontolyticus* (NZ_DS264586.1) organism had the highest coverage (11.3x, 217512 reads) in the HMP staggered dataset, the algorithms on the whole did not perform well on this organism. It was not identified by the Kraken, MetaCV, and MetaPhyler algorithms, and called at a low level by MetaScope (153 reads) (**Figure 2g**) MetaCV mapped the most reads correctly –108,211 (49.7%) and MetaPhlAn was second best, identifying 22,647 (10.4%) of the reads. None of the algorithms identified any of the 2,045 *A. odontolyticus* genes (**Figure 5b**). This poor performance likely results from the fact that *A. odontolyticus* genome annotation in GenBank is incomplete[36]. Conversely, at the species level, five of the six algorithms mapped a high number of reads to *Streptococcus agalactiae* for both the HMP even and HMP staggered datasets (**Figure 2f, 2g**), but only a small number of reads for this organism were present in the truth data. The relative abundance of *Streptococcus mutans* is lower in the algorithm calls as compared to truth, while the relative abundance of *Streptococcus agalactiae* is higher, suggesting that a number of the reads called for *S. agalactiae* are actually from *S. mutans* (**Figure 3b, 3d**). This implies difficulty distinguishing between closely related species. Similarly, a high number of reads are assigned correctly to the *Yersinia* and *Escherichia* genera by Kraken and MetaScope (**Figure 2c**.) However, the algorithms under-assign reads for *Escherichia albertii* and over-assign reads for *Yersinia pseudotuberculosis*, which indicates difficulty in distinguishing between these species (**Figure 2h**).

Overall, algorithms were equally as able to identify organisms in the staggered datasets as in the even datasets, suggesting that accurate read mapping depends more on the database supplied to the algorithm rather than the abundance of the organism in the dataset. Additionally, for the bacterial datasets, Kraken, MetaScope, LMAT, and MetaPhlAn generally agreed on read mapping assignments. However, for the viral datasets, the algorithms missed different sets of organisms – i.e., in **Figure 3i**, LMAT failed to map reads for HIV1, Influenza A virus, Marburg virus, and Machupo virus, whereas MetaScope and Kraken correctly mapped reads for these organisms. However, MetaScope and Kraken both failed to map reads for Human papillomavirus 5, SARS coronavirus, Human papillomavirus 32, and Canine papillomavirus 3, while LMAT succeeded in mapping reads for these organisms. This suggests that for viral datasets, it might be worthwhile to execute both LMAT and one of Kraken or MetaScope, and calculate the union of the results.

The algorithms were also evaluated based on false positive hits (Figure 4). MetaCV and LMAT have diverse error profiles – small numbers of reads are mapped to a high number of false positive organisms. Our past experiences with the MetaScope algorithm suggest that this false positive profile indicates an algorithm has difficulty classifying organisms that are not present in the reference database. Ideally, when an algorithm encounters a novel organism, it should regress up the taxonomic tree until a nearest neighbor for the unknown organism can be established. However, the algorithm may instead report all reference organisms that match the unknown sample to a certain threshold. In contrast, Kraken has a highly concentrated error profiles; fewer than 20 false positive organisms are reported, but several thousand reads are mapped to each of them, suggesting high confidence calls. **Figure 4c and 4d** summarizes the top 20 organisms in terms number of mapped reads, indicating high agreement between Kraken and MetaScope. On the list of false positive genera are several members of the *Enterobacteriaceae* family, including *Shigella, Klebsiella*, and *Enterobacter*. The true positive genera *Salmonella, Escherichia*, and *Yersinia* are members of this family as well. More difficult to explain is the presence of the *Methanolobus* genus, which is a member of the kingdom Archaea and is distantly related to the bacteria in the truth data.

**Figure 4.**
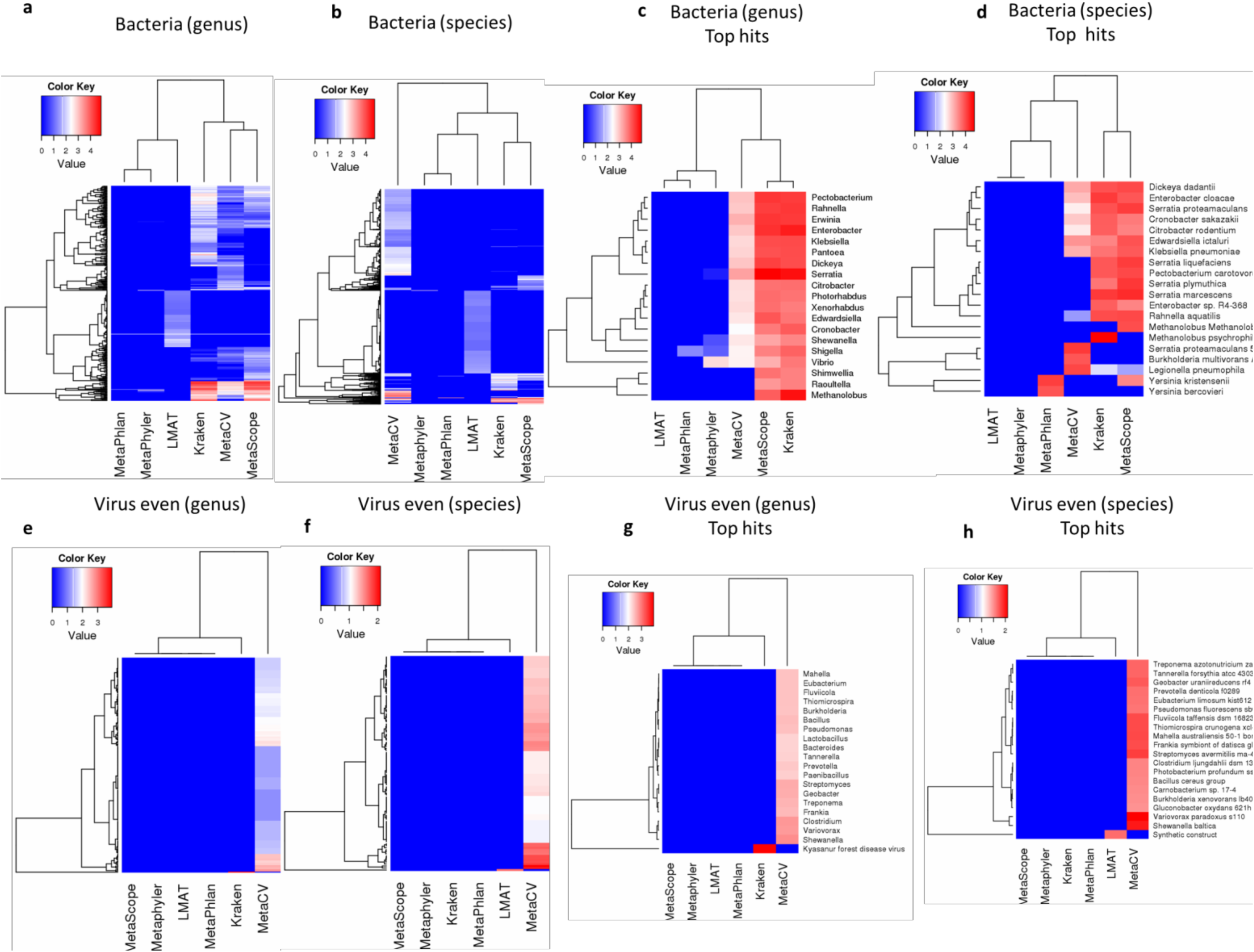
False positive organisms identified to the genus and species level by the 6 metagenomic algorithms. Heatmap color scales are log10 (number of incorrectly assigned false positive reads) for a genus or species. **a**. All false positive genera identified in the bacterial dataset. **b**. All false positive species identified in the bacterial dataset. **c**. 20 false positive genera for the bacterial dataset with the most assigned reads. **d**. 20 false positive species for the bacterial dataset with the most assigned reads. **e**. All false positive genera identified in the virus even dataset. **f**. All false positive species identified in the virus even dataset. **g**. 20 false positive genera for the virus even dataset with the most assigned reads. **h**. 20 false positive species for the virus even dataset with the most assigned reads.

Forthe viral datasets, MetaCV returned a high number of false positives and exhibited poor performance. Kyasanur forest disease virus, a close relative to the true positive Omsk hemorrhagic fever virus was the sole false positive for LMAT, and MetaScope did not report any false positive organisms for either viral dataset.

Finally, the gene calling capabilities of the algorithms were evaluated (Figure 5). Only MetaScope, LMAT, and MetaCV call genes, so these three were included for analysis. For the HMP Even/Staggered, Bacteria, and Virus Staggered datasets, MetaScope identified the most genes correctly out of the three algorithms. LMAT identified more correct genes on the Virus Even dataset (101, compared to 93 for MetaScope).

**Figure 5.**
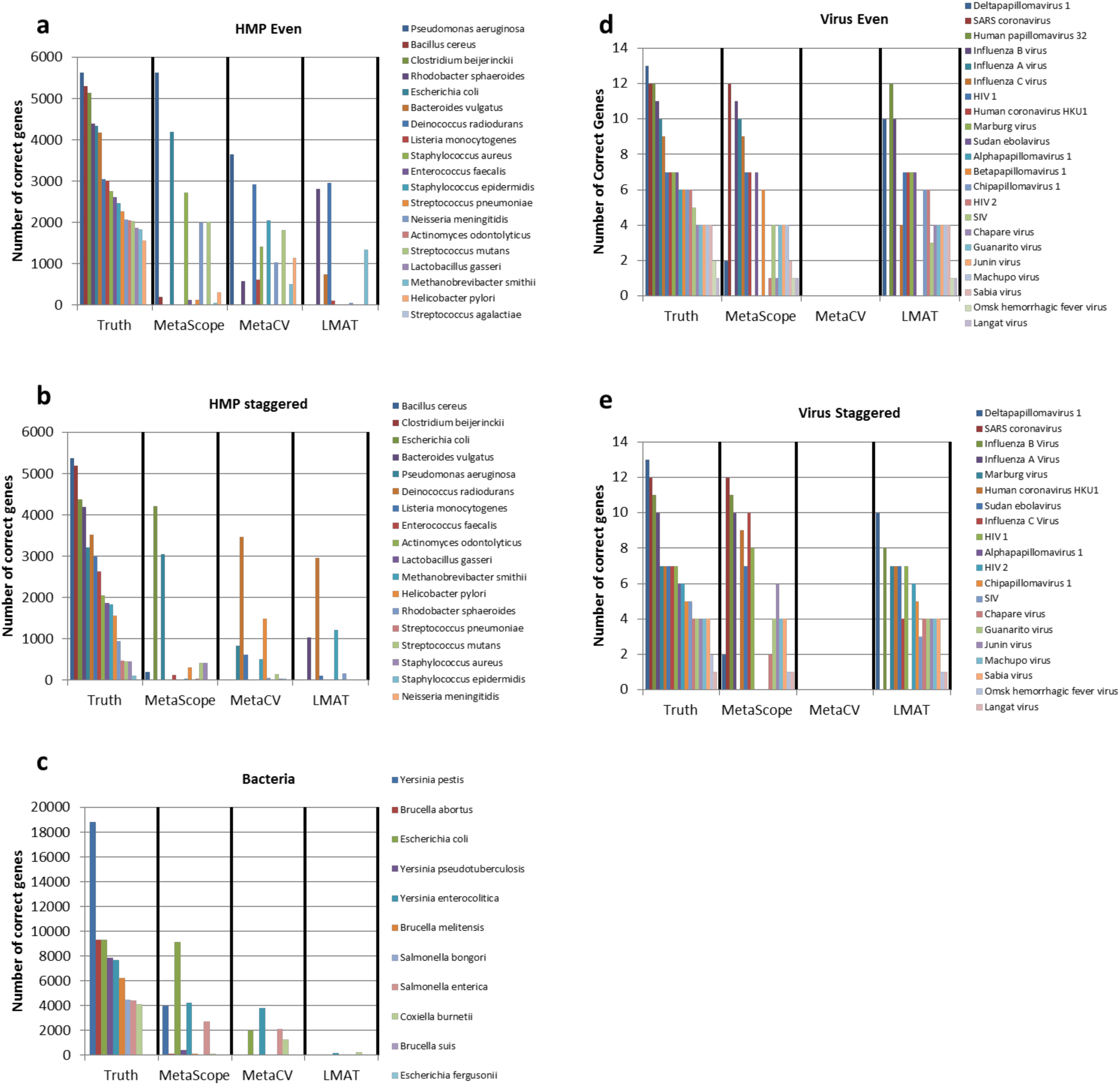
Number of genes correctly identified to the species level across the 5 evaluation datasets (the 6^th^ evaluation dataset consisting of human host reads is not shown). “Truth” column indicates the number of genes with non-zero read coverage in the dataset. MetaScope, MetaCV, and LMAT algorithms provide gene assignment capabilities; Kraken, MetaPhyler, and MetaPhlAn do not call genes and were not included in this evaluation.

## Conclusions

In summary, *in silico* datasets with known truth data for read and gene distribution across different taxons serve as a valuable tool for evaluating algorithm performance. The HMP Even/Staggered, Bacteria, Virus Even/Staggered, and Human datasets generated with FASTQsim elucidate multiple patterns in performance for leading metagenomics algorithms. No algorithm out performed the others in all categories, and the algorithm of choice strongly depends on analysis goals. For bacterial datasets, MetaPhlAn is a clear winner, achieving the lowest runtime, highest ratio of true positives to false positives, and the most precise read mapping. However, MetaPhlAn does not assign genes and does not work on taxons other than Bacteria. LMAT is a clear winner for viral datasets in terms of accuracy, and also provides gene calling functionality. The algorithm most closely matched the relative abundance profile of the truth genera and species across all datasets. However, LMAT also reported the highest rate of false positive genera and species calls on the bacterial datasets. Kraken and MetaScope were the runners up in terms of runtime, ratio of true positives to false positives, and read mapping. MetaScope also performed best for gene mapping, which Kraken does not do. These algorithms performed solidly across all categories evaluated and can be applied most universally across versatile metagenomic applications. MetaPhyler and MetaCV came in last for runtime, ratio of true positives to false positives, and read mapping. They also do not provide results out of the box for viral datasets.

Although viral, bacterial, and human datasets were simulated for this study, the techniques described here can be extended to evaluate metagenomic algorithm performance for other taxa. For example, fungal contamination incidents at medical facilities such as the 2012 incident at the New England Compounding Center[37] can be contained more quickly and effectively with the aid of metagenomic sequencing. Other potential applications include rapid diagnosis of parasite infections[38].

## List of abbreviations

GB: gigabyte
RAM: random-access memory
s: seconds
TB: terabyte
x: fold coverage

## Ethics Committee Approval

Ethics approval was not required for this study because all data was generated *in silico* using references available in GenBank, as indicated in Supplementary Tables 1-3.

## Authors’ contributions

AS implemented FASTQSim updates and generated *in silico* datasets. AS and NC benchmarked algorithm performance on evaluation datasets. AS and DR wrote the manuscript. DR conceived of the study. All authors read and approved the final manuscript.

## Supplementary Materials

**Supplementary Table 1**. Source organisms and coverage levels for HMP Even and HMP Staggered datasets.

**Supplementary Table 2**. Source organisms and coverage levels for Bacterial dataset.

**Supplementary Table 3**. Source organisms and coverage levels for Virus Even and Virus Staggered datasets.

**Figure S1.**
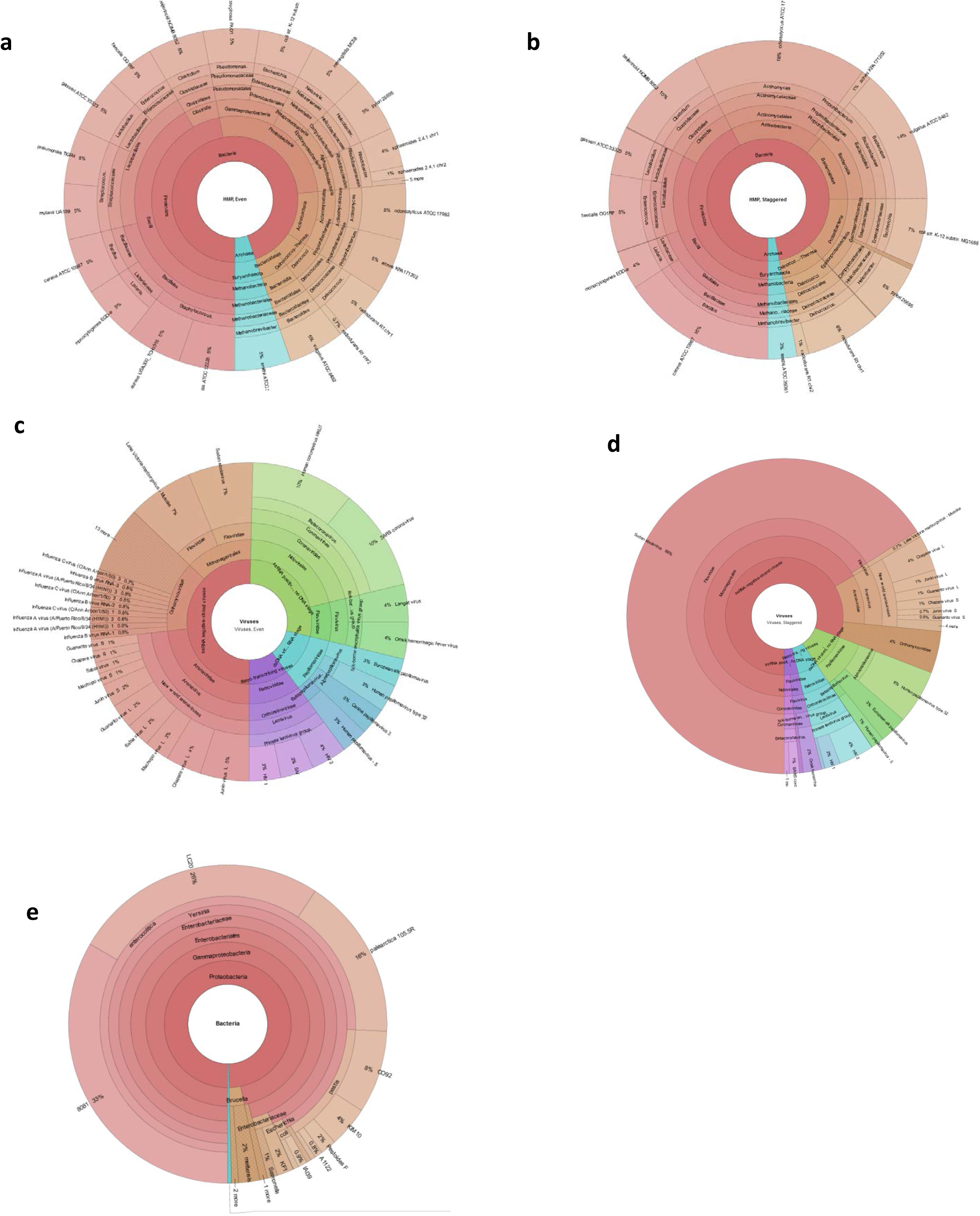
FASTQSim in silico dataset composition to strain level. **a**. 20 bacteria from the Human Microbiome Project (HMP), even coverage levels. **b**. Same 20 bacteria from HMP, staggered coverage levels. **c**. 22 species of viruses across 11 genera, even coverage levels. **d**. Same 22 species of viruses, staggered coverage levels. **e**. 33 strains of bacteria representing 13 species and 5 genera. See Krona HTML files for a-e.

